# Efficacy of *Methylobacterium oryzae* Supplemented *Sargassum wightii* Seaweed Liquid Fertilizer on Chilly and Tomato Plant Growth

**DOI:** 10.1101/2020.03.16.994640

**Authors:** Athiappan Murugan, Anandan Rubavathi, Kannan Visali, Vijayasingh Neginah

## Abstract

Liquid seaweed fertilizer gaining interest due to their use in crop productivity. The present study was designed to evaluate the impact of *Methylobacterium oryzae* amended liquid seaweed fertilizer on domestic plants like chilly and tomato. The *Sargassum wightii* methanolic extract was maximum (137 mg/g) among other solvent tested. Extract showed 0.456mg/mL, 1.587 mg/mL, 0.78 mg/g, 5.27 mg/g, 0.63 mg/mL, 0.98 mg/mL, 0.285 g/mL and 0.546 μg/mL of total phenolic, total flavonoids, total chlorophyll, total carotenoids, total protein, total carbohydrates, total lipids and total aminoacids respectively. The maximum survival of *Methylobacterium oryzae* was observed at 40 % of *Sargassum wightii* SLF extract with 3% *Methylobacterium oryzae* culture concentration and has viability of 750 × 10^6^ CFU/l after 6 months. Impact of foliar sprayed liquid fertilizer on the chlorophyll, internode and shoot length had been promising over seed soaked.

## Introduction

Due to population rise and to fulfill the food demand, in order to meet the increasing demand chemical fertilizers are used (Parr et al. 1994). The application of chemical fertilizers leads to soil pollution and health hazards to the living organisms. Chemical fertilizers are quite expensive and this leads to the rise in cost of production. To meet the increasing demand many viable options are available, among this one of such options is seaweed as a fertilizer. The use of seaweed as manure in farming practice is very ancient and was prevalent among the Romans (Galbiattia et al. 2007). The use of seaweeds as bio-fertilizers in horticulture and agriculture has increased in the recent years (Hong et al. 2007). Seaweed liquid fertilizer (SLF) of *Sargassum* species contains macronutrients, trace elements, organic substance like amino acids and plant growth regulators.

For sustainable crop production, plant growth promoting symbiotic and non-symbiotic free-living bacteria are employed as external source of nitrogen (Lugtenberg et al 2009). *Methylobacterium suomiense* CBMB120, a plant growth promoting (PGP) root and shoot colonizer has been used widely (Kennedy et al. 2004). Many research studies have shown the beneficial effect of seaweeds extracts in stimulating growth of plants (Blunden et al. 1993)as it possesses plant nutrients and hormones such as auxins and gibberellins (Wu et al. 1997). Several research reports are available on the beneficial effects of seaweed extractsas natural regulators and osmoregulants (Featonby-Smith and VanStaden 1983). *Methylobacterium* sp. is known for their eco-friendly ability to support plant growth. They can able to grow on a wide range of multi-carbon substrate such as methanol, formate, and formaldehyde to produce plant growth promoting metabolites (Holland 1997). *Methylobacterium* sp. have the potential to fix the atmospheric nitrogen in the non-leguminous plants (Galbiattia et al. 2007) and (Miguel et al. 2007).

*Methylobacterium* sp. have been reported to have many plant growth promoting abilities including nitrogen fixation, phosphate solubilization, production of plant growth promoters, and biological disease control (Holland and Polacco 1992; Munusamy et al. 2004; and Omer et al. 2004). The metabolites present in the *Methylobacterium* sp. have the potency to stimulate seed germination and plant growth (Bhagirath and David 2010). However, not much is known about the influence of *Methylobacterium* sp. with seaweed liquid fertilizers on plant growth. With this limitation, the present study was aimed to study the influence of *Methylobacterium oryzae* inoculum along with liquid fertilizers in the plant growth promoting effects on *Lycopersicon esculentum* L (Tomato) and *Capsicum annuum*L (Red pepper).

## Materials and Methods

### Collection of Seaweeds

The brown seaweeds *Sargassum wightii* were collected from the coastal region, Palladam (10.9997° N, 77.2807° E), Rameshwaram (9.2876° N, 79.3129° E), Tamil Nadu, India. The seaweed was cleaned with distilled water and then immediately placed in a freezer (−40 °C) and freeze-dried for 24 hrs. The dried sample was then ground to powder and stored in a sealed bag at −40 °C for further analysis.

### Preparation of Extracts

The freeze-dried seaweed powder was soaked with various organic solvents (1:20, w/v) namely ethanol (80 %, v/v), methanol (80%, v/v), chloroform (80 %, v/v), ethyl acetate (80%, v/v) and petroleum ether and water at room temperature for 24 h in a shaking incubator to extract plant growth promoting components (Airanthi et al. 2011). The slurry was filtered through Whatman No.1 filter paper and filtrates were collected in a container. The residues were re-soaked again under the same conditions. The combined filtrate was evaporated in a rotary evaporator under vacuum at 40 °C to obtain the dried extract.

### Optimization of Seaweed Extraction

To know the extraction abilities of various solvents, 10 g of dried seaweed powder was weighed and soaked with 200 ml of different solvents as mentioned above. To optimize the specific ratio of solvent to seaweed various concentration ratio of seaweed and methanol (w/v) of 1:250, 2:250, 3:250, 4:250 and 5:250 were employed for extraction.

### Physicochemical Analysis of Seaweed Extract

#### Total Phenolic Content (TPC)

The Total Phenolic Content (TPC)(Patricia et al. 2009), Total Flavonoids Content (TFC) (Jim et al. 2002), Total Carotenoids Content (TCC), Total Chlorophyll Content (TCPC), Total Protein (TP), Total Carbohydrate (TC), Total Lipids (TL) and Total aminoacids (TAA) of *Sargassum wightii* methanolic extracts were estimated using the standard methods (Kirk and Allen 1965; Samuel 1964).

### Supplementation of *Methylobacterium* sp. with Seaweed Extract

The bacterial culture of *Methylobacterium oryzae* with varied concentration viz., 0.5%, 1%, 1.5%, 2.5% and 3% were inoculated with different concentration of *Sargassum wightii* extract (5%, 10%, 15%, 20%, 25%, 30%, 35%, 40%, 45% and 50%). Among the tested combination, the combination showed with higher growth was taken for culture viability study and was evaluated at 15 days interval for 6 months. After incubation, bacterial growth was recorded by the colony forming a unit and the mean of log CFU expressed from five random samplings.

To evaluate the impact of *Sargassum wightii* seaweed liquid fertilizers (SLF) supplemented with *Methylobacterium oryzae* combination on seed germination and vegetative growth of *Lycopersicon esculentum* L (Tomato) and *Capsicum annuum* L (Red pepper) four different treatments were performed viz.,control, *Sargassum wightii* SLF extract, *Methylobacterium oryzae*, *Sargassum wightii* SLF extract with *Methylobacterium oryzae* 0.5% of *Sargassum wightii* SLF was prepared and the sowing seeds (Tomato and Chilli) were soaked for 24 hrs. The seeds were sowed and observed for germination and early growth.The soil sterilized in the autoclave for 121°C for 1 h and was used for the pot study experiment. *Sargassum wightii* SLF extract with *Methylobacterium oryzae* was prepared with water and was sprayed at 10 days interval using hand sprayers after the germination of seeds.

### Enumeration of Rhizosphere and Phyllosphere Bacterial Population

Owing to evaluate the impact of bacterial inoculum in 10g of pot soil sample was added into 100ml of sterile distilled water and kept in shaking incubator at 200rpm for 20 mins. Serial dilution of the potting soil was prepared and plated on nutrient agar using pour plate method. The soil solution seeded plates were incubated at 37 °C for 24 h. Rhizosphere bacterial population numbers were expressed as mean values of ten different samples.

The leaves of Tomato and Red pepper were detached aseptically from the branches of experimental plants and washed separately. Specific leaves are kept in a shaker for one hour with 100 ml sterile distilled water to isolate the bacteria and the suspension was treated as stock. For enumeration of the total viable count,serial dilutions of the stock solution were prepared up to 10^−5^ and an aliquot of 100μl from each the dilutions were plated separately in nutrient agar media. Phyllosphere bacterial population counts were recorded bya mean value of ten different samples.

### Plant Growth Index

The Plant growth index is used as a quantitative indicator of plant growth rate and to compare the size of the plants grown under different systems. Values are expressed in mean of ten plants analyzed for each treatment. Plants shoot heights (aerial parts only), number of nodes and length of the internodes were measured on the 90^th^ day.

### Analysis of Plant Chlorophyll

The leaves were collected from the potted plants of Tomato and Red pepper and chlorophyll was extracted with dimethyl sulphoxide (Hiscox and Israelsta 1979). For the chlorophyll extractions, glass centrifuge vials containing 7 ml DMSO was preheated to 65°C in a water bath. In the preheated vials, three leaves discs (each 3.038 cm^2^; approx. 100 mg. wt. total) were incubated for 20 min. The DMSO was used as blank and absorbance of *Sargassum wightii* SLF extract with *Methylobacterium oryzae* treated plant leaves was measured at 645 and 663 nm after 30 minutes. Arnon’s (1949) equation was employed to calculate the chlorophyll content. Chla (g l-^1^) = 0.0127 A663 – 0.00269 A645; Chlb (g l-^1^) = 0.0229 A645 – 0.00468 A663; Total chlorophyll (g l-^1^) = 0.0202 A645 + 0.00802 A663. The chlorophyll concentration of the extract calculated from these equations was then converted to leaf chlorophyll content (mg Chl cm-^2^ leaf area).

## Results

The extractive values of the *Sargassumm wightii* were as follows methanol (137 mg/g) > water (94 mg/g) > ethanol (80 mg/g > chloroform 20 mg/g > petroleum ether (10mg/g). Among the various ratio tested, 40:250 ratio (Seaweed: Methanol) yielded highest frequency (14.93%; 5.972g/L) extracts.

### Physicochemical Analysis of *Sargassum* Species

#### Total Phenolic Content (TPC)

The physico-chemical analysis of *Sargassum wightii* revealed that the varied number of metabolites. The *Sargassum wightii* methanolic extracts showed 0.456mg/mL, 1.587 mg/mL, 0.78 mg/g, 5.27 mg/g, 0.63 mg/mL, 0.98 mg/mL, 0.285 g/mL and 0.546 μg/mL of total phenolic, total flavonoids, total chlorophyll, total carotenoids, total protein, total carbohydrates, total lipids and total aminoacidsrespectively^25^.

### Supplementation of Seaweed Extract with *Methylobacterium oryzae*

The survival rates of *Methylobacterium oryzae* at different concentrations of *Sargassum wightii* SLF extract were analyzed. The maximum survival of *Methylobacterium oryzae* was observed at 40 % of *Sargassum wightii* SLF extract with 3% *Methylobacterium oryzae* culture concentration after 6 months of viability was 750 × 10^6^ CFU (Fig. 01). In this study, the *Sargassum wightii* SLF is used as carrier material which showed the viability of *Methylobacterium oryzae* for about six months. After 6 months, the *Methylobacterium oryzae* showed a decline in growth.

**Fig. 01.**
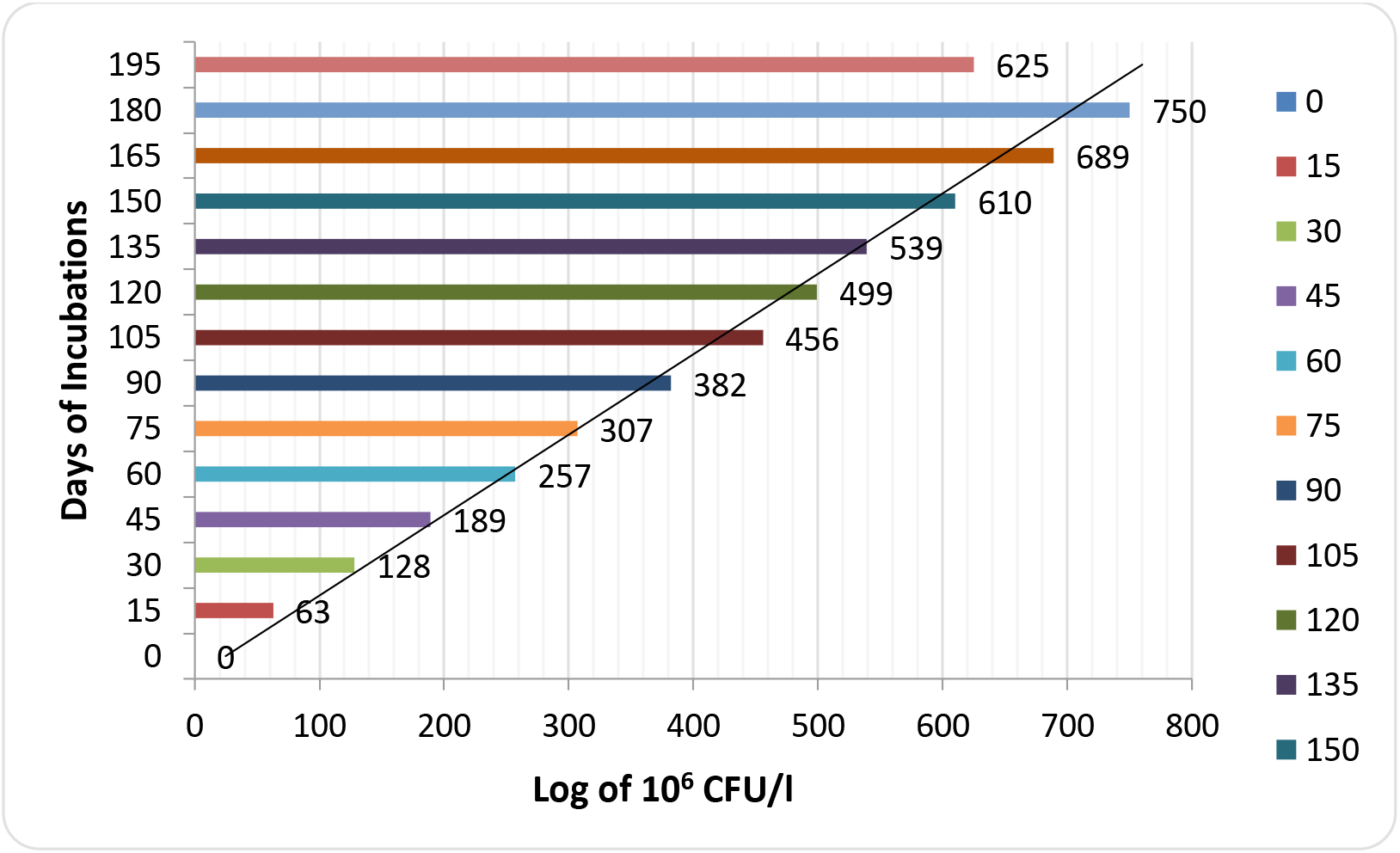
Enumeration of Rhizosphere and Phyllosphere flora

A load of *Methylobacterium oryzae*in the rhizosphere of tomato and red pepper was increased in the experimental conditions viz., seeds soak and foliar spray conditions (Fig. 02). Similar trend like rhizosphere population in the experimental conditions, Phyllosphere flora also increased with a period of 3 months in the pot treated with *Sargassum wightii* SLF extract with *Methylobacterium oryzae* (Fig. 03).

**Fig. 02.**
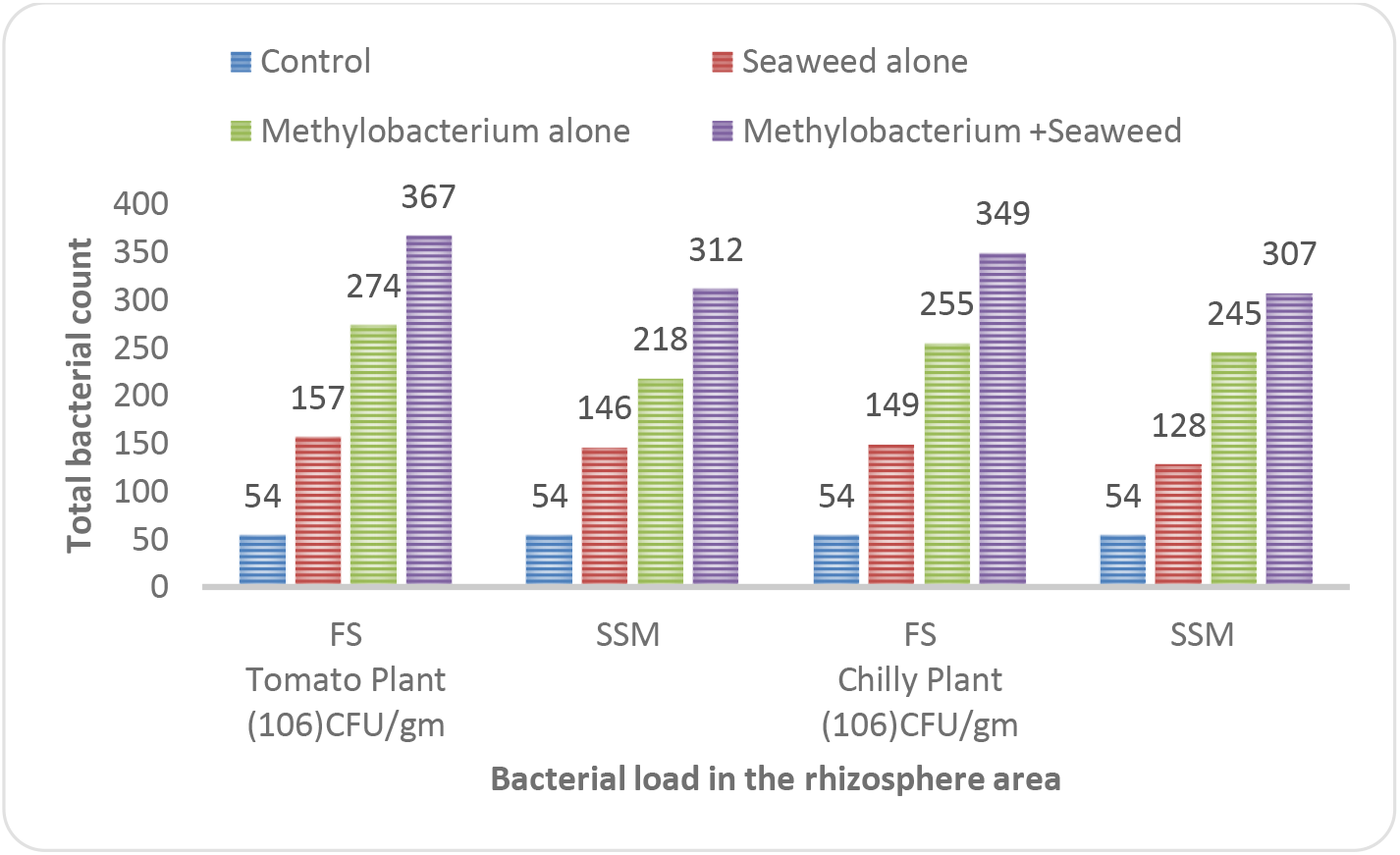
Bacterial load of *Methylobacterium oryzaein* the rhizosphere of tomato and red pepper

**FIG. 03.**
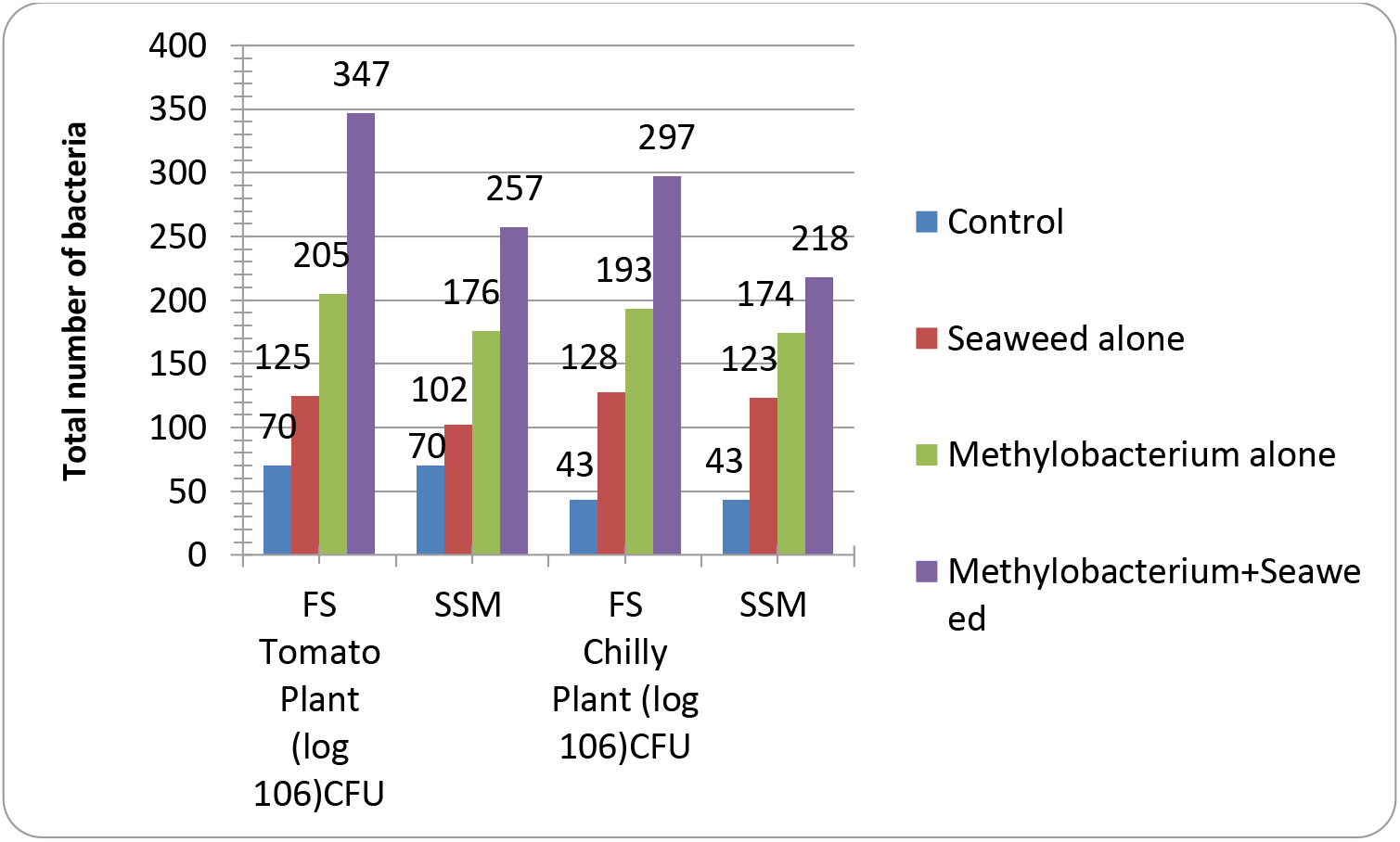
Phyllosphere flora in the pot plant treated with *Sargassum wightii* SLF extract amended with *Methylobacterium oryzae*

### Plant Growth Index

Foliar spray of *Sargassum wightii* SLF extract with *Methylobacterium oryzae* combination treated red pepper and tomato showed gradual increase in plant shoot growth than foliar application of *Sargassum wightii* SLF extracts and *Methylobacterium oryzae* alone. This study results suggested that *Sargassum wightii* SLF extract with *Methylobacterium oryzae* combination considered as good fertilizer. The flowering of tomato and red pepper was observed after 90^th^days.

Foliar spray of *Sargassum wightii* SLF extract with *Methylobacterium oryzae* combination treated tomato and red pepper plant displayed 49.9 cm and 33.5 cm of shoot length with 3.7 node and 3.9 and 2.8 cm and 1.1 cm inter-node length respectively (Fig. 04). Similarly, the chlorophyll contents of the tomato and red pepper were increased with the foliar application of *Sargassum wightii* SLF extract with *Methylobacterium oryzae* combination. The plant growth was higher in the foliar application of *Sargassum wightii* SLF extract with *Methylobacterium oryzae* combination than seed-soaked method (Table 6; Fig 04–06).

**FIG. 04.**
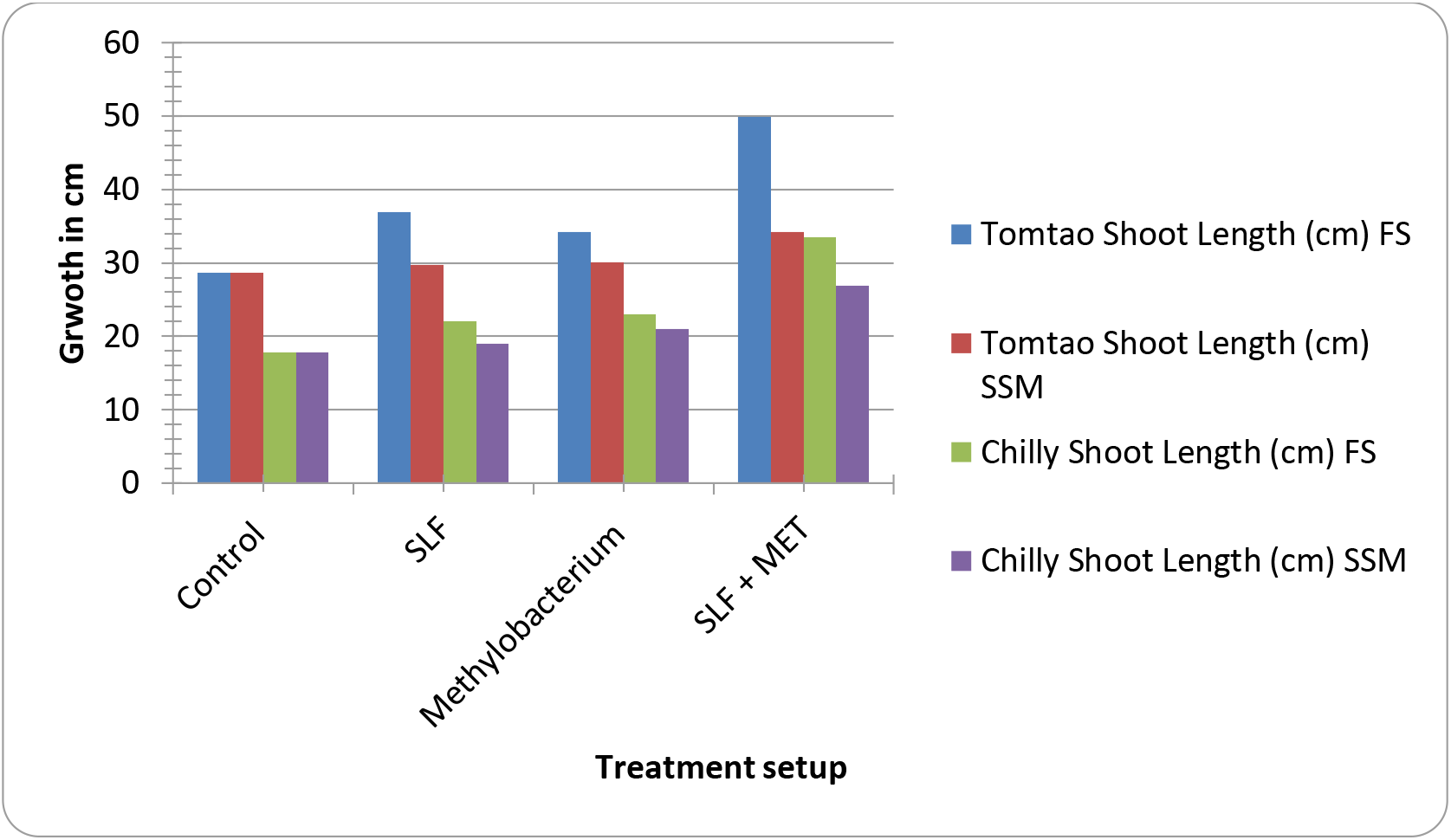
Foliar spray of *Sargassum wightii* SLF extract with *Methylobacterium oryzae* combination treated tomato and red pepper plant

**FIG. 05.**
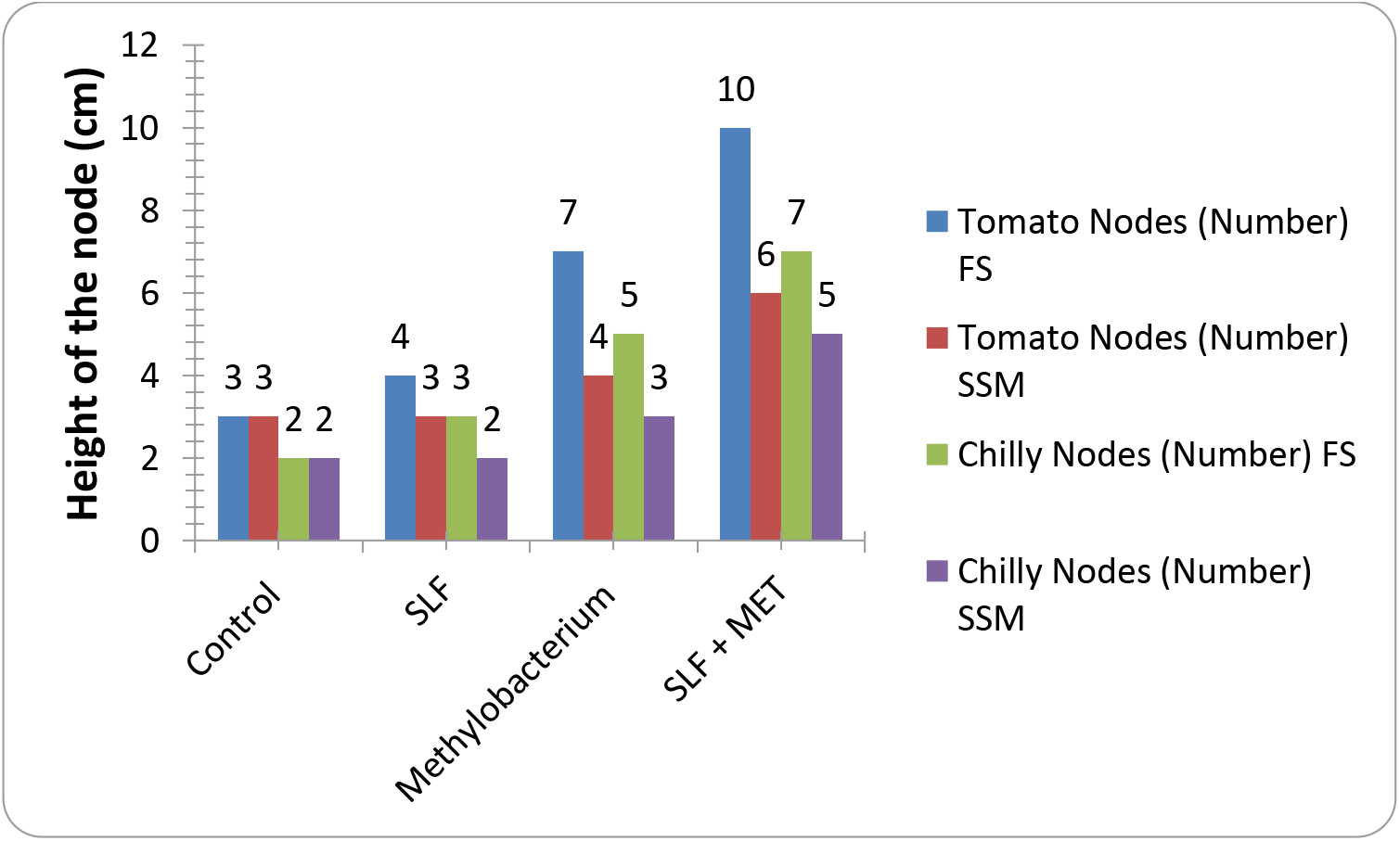
Effect of liquid fertilizer applied by spray and seed socking on the plant growth (node)

**FIG. 06.**
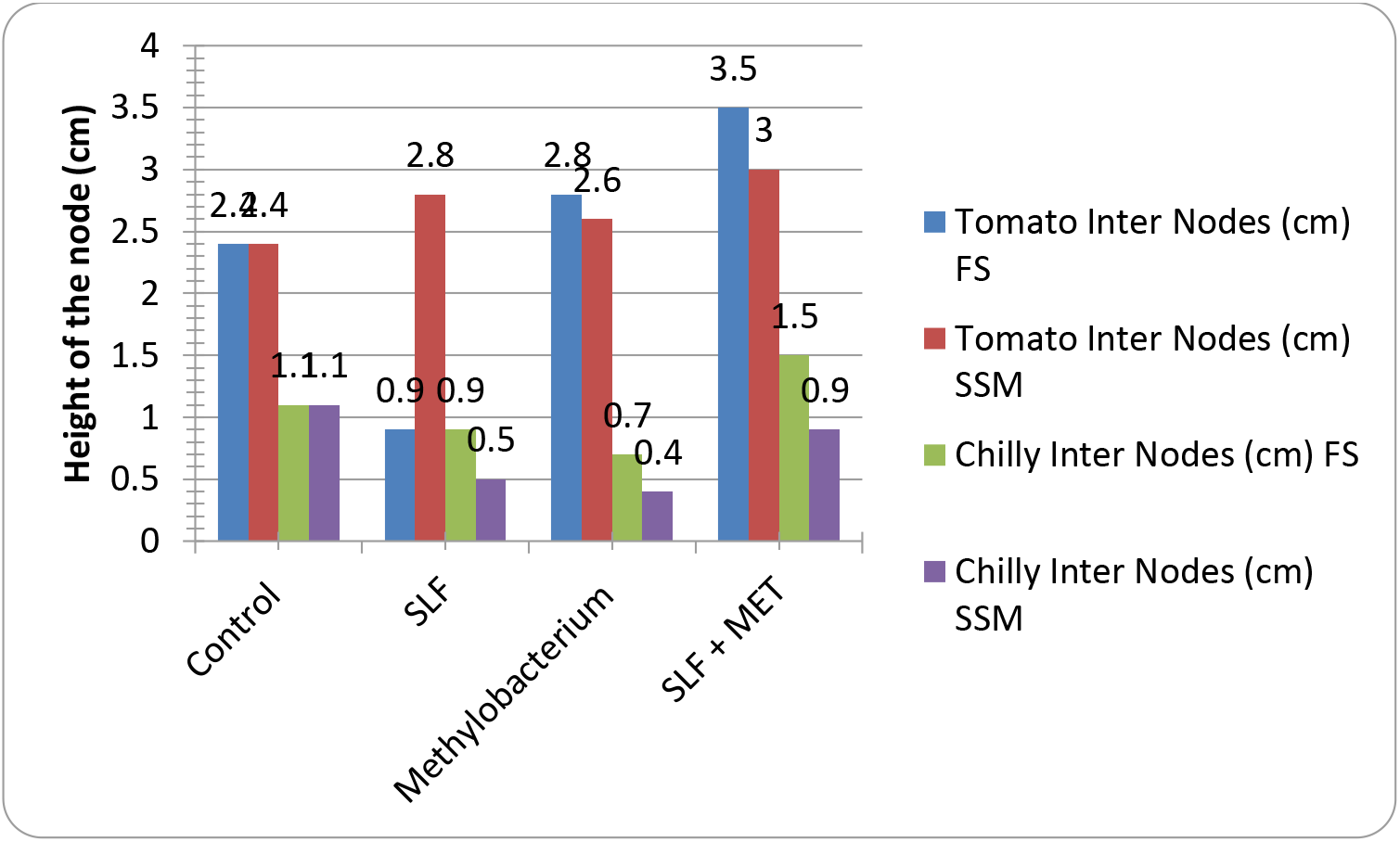
Effect of liquid fertilizer applied by spray and seed socking on the plant growth (internode)

## Discussion

Seaweed liquid fertilizer is a new generation of natural organic fertilizers containing highly effective, nutritious, promotes seed germination, increase yield and resistant ability of many crops. Modern agricultural practices need more amount of fertilizers for higher yield to satisfy human requirements. The seaweed extracts contain plant growth regulators (auxins, gibberellins and cytokinin), growth promoters, carbohydrates, amino acids, antibiotics, and vitamins consequently which enhance the yield and quality which induce the yield of crops, seed germination, resistance to frost, fungal and insect attacks (Erulan et al.1824).

Variations in the seaweed extractive values are attributed to polarities of different solvents employed for the extraction. Based on the metabolites in the plants, the influence of SLF on the plant growth found varied. Previous studies on growth promoting effect of various SLF indicated that concentration and metabolites influence plant growth. In the present study, extractive percentage was high in the methanol than other studied solvents (Matanjun et al. 2008). The phytochemical and biochemical analysis also confirmed the occurrence of more amount metabolites in the methanolic extracts and hence the methanolic extracts of *Sargassum wightii* (*Sargassum wightii* SLF) chooses for further analysis.

Similar to the present study, high extractive values were obtained in the methanolic extracts derived SLF of *Sargassum polycystum* (4.05%), *A. spicier* (5.01%), *G. edulis* (3.98%) and *E. kappaphycus* of about 2.85% (Bhagirath and David 2010; Kuda et al. 2005) observed the increased organic fertilizer application level with inoculation of *M. oryzae* CBMB20 significantly improved red pepper plant growth. The application of *M. extorquens* MM2 substantially increased the seed germination percentage and seedling growth of tomato. In the present study, the foliar application of *Sargassum wightii* SLF extract with *Methylobacterium oryzae* combination also increased the vegetative growth of red pepper and tomato^16^ also observed the significant increment in the accumulation of IAA and cytokinin in the red pepper treated with *Methylobacterium* sp. CBMB20. Munusamy et al. (2004) studied the co-inoculation effect of *M. Oryzae* CBMB20 with nitrogen-fixing *Azospirillum brasilense* CW903 or a phosphate solubilising bacterium *Burkholderia pyrrocinia* CBPB-HOD had positive effect on tomato, red pepper and rice crop. Application of seaweeds and seaweed extracts triggers the growth of beneficial soil microbes and secretion of soil conditioning substances by these microbes.

Seaweeds and seaweed products enhance plant chlorophyll content (Blunden et al. 1993). Application of a low concentration of *Ascophyllum nodosum* extract on foliage of tomatoes produced leaves with higher chlorophyll content than those of untreated controls. This increase of chlorophyll content is due to betaines in the seaweed extract which reduces chlorophyll degradation. Glycine betaine delays the loss of photosynthetic activity by inhibiting chlorophyll degradation during storage conditions in isolated chloroplasts. An observation was made in *Scytonema* sp., *Vigna mungo* and in *Vignasinensis* (Sivasankari. 2006). The present study also suggests the foliar application of *Sargassum wightii* SLF extract with *Methylobacterium oryzae* combination increased the chlorophyll content of tomato and red pepper.

## Conclusion

Seaweed liquid fertilizers extracted from *Sargassum wightii* had influenced agricultural productivity. The physico-chemical analysis has shown that these extracts contain various factors such as cytokinins, gibberellins, trace elements, vitamins, amino acids, antibiotics and micronutrients. Plant growth promotion has been pertinent to influential effect of *Sargassum wightii* SLF extract. Further *Methylobacterium oryzae* also influence positively the plant growth and that has been observed in root length, shoot length, fresh weight and dry weight. Hence, it can be recommended to supplement SLF with plant growth promoting bacteria for greater benefits.

## Acknowledgements

Both the authors acknowledge the DST-FIST facilities (Ref No. SR/FST/LSI-640/2015C).

## Author contribution

Athiappan Murugan: Original Concept

Anandan Rubavathi: Culture experiment

Kannan Visali: Manuscript editing

Vijayasingh Neginah: Data analysis

